# Loss of SUR1 subtype K_ATP_ channels alters antinociception and locomotor activity after opioid administration

**DOI:** 10.1101/2020.12.02.408450

**Authors:** Gerald Sakamaki, Kayla Johnson, Megan Mensinger, Eindray Hmu, Amanda H. Klein

## Abstract

**Background:** Opioid signaling can occur through several downstream mediators and influence analgesia as well as reward mechanisms in the nervous system. K_ATP_ channels are downstream targets of the μ opioid receptor and contribute to morphine-induced antinociception.

**Aims:** The aim of the present work was to assess the role of SUR1-subtype K_ATP_ channels in antinocicpetion and hyperlocomotion of synthetic and semi-synthetic opioids.

**Methods:** Adult male and female mice wild-type (WT) and SUR1 deficient (KO) mice were assessed for mechanical and thermal antinociception after administration of either buprenorphine, fentanyl, or DAMGO. Potassium flux was assessed in the dorsal root ganglia and superficial dorsal horn cells in WT and KO mice. Hyperlocomotion was also assessed in WT and KO animals after buprenorphine, fentanyl, or DAMGO administration.

**Results:** SUR1 KO mice had attenuated mechanical antinociception after systemic administration of buprenorphine, fentanyl, and DAMGO. Potassium flux was also attenuated in the dorsal root ganglia and spinal cord cells after acute administration of buprenorphine and fentanyl. Hyperlocomotion after administration of morphine and buprenorphine was potentiated in SUR1 KO mice, but was not seen after administration of fentanyl or DAMGO.

**Conclusions:** These results suggest SUR1-subtype K_ATP_ channels mediate the antinociceptive response of several classes of opioids (alkaloid and synthetic/semi-synthetic), but may not contribute to the “drug-seeking” behaviors of all classes of opioids.

## Introduction

Classical opioid analgesia works through G-protein coupled receptor (GPCR) signaling, followed by a decrease in Ca^2+^ activity and an increase in K^+^ signaling through direct or indirect Gαi-protein modulation (Al-Hasani and Bruchas, 2011). Activation of potassium channels such as inwardly-rectifying potassium channels (K_ir_) and G-protein coupled inwardly-rectifying potassium channels (GIRKs) are suggested as potential targets for broadening analgesic therapies through ligand bias at opioid receptors (Clayton et al., 2009; Kliewer et al., 2019). Previous studies indicate buprenorphine and DAMGO signal through K_ir_ (Knapman et al., 2014), but other synthetic opioids such as fentanyl may not share the same signaling pathways (Kanbara et al., 2014). Research into various signal transduction pathways and targets of opioids are ongoing in search of novel targets for pain relief. For example, TRV130 and PZM21, were developed to specifically target the μ opioid receptor and G-protein activation without recruiting β-arrestin signal transduction pathways with the purpose of decreasing adverse effects of these opioids (DeWire et al., 2013; Hill et al., 2018; Manglik et al., 2016; Soergel et al., 2014).

Among the K_ir_s, ATP sensitive potassium channels (K_ATP_) are another downstream target of the μ opioid receptor (Cunha et al., 2010) and produce analgesia in rodent pain models (Chi et al., 2007; Luu et al., 2019a; Niu et al., 2011; Wu et al., 2011a). Previous studies indicate activation of K_ATP_ channels potentiates morphine analgesia (Afify et al., 2013), while loss of K_ATP_ channel subunit expression decreases morphine antinociception(Fisher et al., 2019). K_ATP_ channels are found in the peripheral nervous system, spinal cord, and brain (Babenko et al., 1998; Seino and Miki, 2003) and form an octamer which consists of four Kir6.X subunits and four regulatory sulfonylurea (SUR) subunits (Ashcroft, 2005; Babenko et al., 1998). Kir6.X and SURX are also expressed in pairs, with different combinations being more prominent in certain tissues, such as Kir6.2/SUR1 in the peripheral nervous system (Babenko et al., 1998; Zoga et al., 2010), Kir6.1/SUR1 and Kir6.1/SUR2 in the spinal cord (Wu et al., 2011b), and various combinations of K_ATP_ channel subunits across the brain depending on region.

The administration of drugs of abuse, including opioids, increase locomotor activity in rodents, which is thought to be related to drug seeking behaviors. Behavioral studies indicate that modulation of K_ATP_ channel activity affects these behaviors in rodents. Iptakalim, a K_ATP_ channel opener, was found to attenuate nicotine-induced dopamine and glutamate release in the nucleus accumbens of rats (Liu et al., 2006; Volf et al., 2012), and systemic administration of the K_ATP_ channel agonists iptakalim, cromakalim, and pinacidil decreases amphetamine induced hyperlocomotion(Rosenzweig-Lipson et al., 1997; Sun et al., 2010). These data suggest activation of K_ATP_ channels in the central nervous system can also attenuate drug seeking behaviors in addition to nociception. Whether an attenuation of antinociception and a potentiation of hyperlocomotion are found after administration of synthetic and semi-synthetic opioids after loss of K_ATP_ channel activity is not known at this time.

In this study, opioid-induced behaviors were measured using mechanical and thermal sensitivity and open field testing, to investigate antinociceptive and locomotor effects of buprenorphine, DAMGO, and fentanyl in SUR1-deficient mice, respectively. Attenuation of antinociception after buprenorphine, DAMGO, and fentanyl administration was found in SUR1 knock out (KO) compared to wildtype (WT) mice, but hyperlocomotion was only largely potentiated after morphine or buprenorphine administration. These data suggest several opioids can stimulate K_ATP_ channel activation to induce analgesia, but K_ATP_ channel activity with regards to drug seeking or other opioid-induced behaviors may be dependent on the opioid ligand used, possibly due to differences in downstream signaling cascades across the nervous system.

## Materials and Methods

### Experimental models

All experimental procedures involving animals were approved and performed in accordance with the University of Minnesota Institutional Animal Care and Use Committee. Breeding pairs of SUR1-deficient mice (SUR1 KO) and SUR1^flx/flx^ (SUR1 flox) mice were obtained from the laboratory of Dr. Joseph Bryan at the Pacific Northwest Research Institute (Seattle, WA, United States) as used in previous studies (Nakamura and Bryan, 2014; Seghers et al., 2000) and kept on a C57Bl6N background. SUR1 WT littermates were used as controls to SUR1 KO mice. Behavioral experiments were performed on adult male and female mice (>5 weeks of age, 20-40 g) and tissue isolation was performed on adult mice (4-8 weeks, 20-40 g). Genotype was verified by PCR as in previous studies (Luu et al., 2019a; Seghers et al., 2000).

### Drugs

Buprenorphine (5.83 mg/kg; PHR172, Sigma Aldrich, St. Louis, MO), DAMGO (10 mg/kg; Bachem, Bubendorf, Switzerland), morphine (5 and 15 mg/kg; Spectrum Chemical, New Brunswick, NJ), and fentanyl (0.25 mg/kg; F2886, Sigma Aldrich) were diluted in saline (vehicle) and administered through 100 μL subcutaneous injections. The drug concentrations used on this study were found in previous studies to elicit significant antinociceptive affects in mice (Cowan et al., 1977; Lutfy et al., 2003; Martucci et al., 2004). Some mice were used in multiple studies. To avoid lingering effects from previous drug injections and tests, a minimum of three days elapsed between each injection and test. Drug and test order were randomized.

### Mechanical Paw Withdrawal

Mice were acclimated on several separate occasions in individual acrylic containers on a shared wire mesh floor one week prior to the start of experimentation. Mechanical paw withdrawal thresholds were measured using electronic von Frey testing equipment (Electric von Frey Anesthesiometer, 2390, Almemo^®^ 2450, IITC Life Science, Woodland Hills, CA) (Martinov et al., 2013). Thresholds were measured as the amount of force in grams required to elicit a nocifensive response on the plantar surface of both rear hind paws. Baseline thresholds consisted of 3-5 measurements from each hind paw prior to drug administration and were averaged. Post-drug administration thresholds from both hind paws were measured at 3, 15, 30, 45, 60, and 120 minutes post-injection.

### Thermal Paw Withdrawal

Mice were acclimated on multiple separate occasions to individual acrylic containers on a shared glass floor heated to 30°C one week prior to the start of each experiment. Thermal paw withdrawal latencies were measured using a modified Hargreaves Method (Plantar Test Analgesia Meter, 400, IITC, Woodland Hills, CA) (Banik and Kabadi, 2013). Latencies were measured as the amount of time in seconds for a heat radiant light beam focused on the plantar surface of each hind paw required to elicit a nocifensive response (Cheah et al., 2017). A time limit of 20 seconds of exposure to the beam was implemented to avoid tissue damage. Mice were restrained using laboratory-made restrainers (Machholz et al., 2012). Baseline latencies consisted of 3 measurements per hind paw prior to drug administration and were averaged. Post-drug administration latencies from both hind paws were measured at 3, 15, 30, 45, 60, and 120 minutes post-injection.

### Primary Culture of DRG and Spinal Cord Dorsal Horn

Animals were anaesthetized with 5% isoflurane in oxygen and decapitated prior to tissue isolation. DRG isolation was similar to previous reports but are summarized below (Fisher et al., 2019). DRG were extracted from male SUR1 WT and SUR1 KO mice (4-8 weeks, 20–40 g) and placed on ice in Petri dishes containing 1X Hank’s Balanced Salt Solution (HBSS, SH30588.02, GE Healthcare Life Sciences, US). Ganglia were minced using surgical micro scissors and incubated in a papain solution (32 μL papain (27.3 U/mg, no. 3126; Worthington Biochemical, Lakewood, NJ, US and 1 mg l-cysteine, Sigma Aldrich) in 1.5 mL 1X HBSS in a 37°C rocking water bath for 10 minutes. The samples were then centrifuged at 1600 rpm for 2 min (Eppendorf 5415R, 200xg) and the supernatant aspirated before the tissue was incubated in a collagenase solution containing 2 mg/mL collagenase type II (CLS2; Worthington Biochemical) in 5 mL 1X HBSS in a 37°C rocking water bath for 10 minutes. Cells were centrifuged at 1500 rpm for 2 minutes (Beckman Allegra X15R, 524xg), supernatant was removed and tissue was carefully triturated using fire-polished glass pipettes with 2 mL pre-warmed media. The media contained Eagle’s Minimum Essential Medium with Earle’s salts and L-glutamine (10-010-CV, Corning^®^, Corning, NY, US), 10% v/v horse serum (26050-088; Gibco, Thermo Fisher Scientific, Waltham, MA, United States), 1% v/v 100X MEM vitamins solution (11120-052; Gibco, Thermo Fisher Scientific, Waltham, MA, United States) and 1% v/v penicillin-streptomycin (15140-122; Gibco, Thermo Fisher Scientific, Waltham, MA, United States). DRG were then centrifuged for 2 min at 1000 rpm (Beckman Allegra X15R, 233xg), supernatant aspirated, and the pellet resuspended in 5 mL pre-warmed media. Isolated DRG were plated in 100 μL aliquots into clear bottom, black-walled 96-well plates coated with 2.5 μg/cm^2^ poly-D-lysine and incubated in a humidified incubator at 37°C with 5% CO2 (~1.5×10^5^ cells/well).

Dorsal horn cell isolation was modified from a previous report(Cao et al., 2017). The spinal cord was isolated from male SUR1 WT and SUR1 KO mice (4-8 weeks, 20–40 g) and placed on ice in Petri dishes containing 10 mM HEPES (BP310-1, Thermo Fisher Scientific, Waltham, MA, United States) in 1X Hank’s Balanced Salt Solution (SH30588.02, GE Healthcare Life Sciences, United States). The dura was removed and the spinal cord was bisected lengthwise. The halves were bisected again and a quarter width of the spinal cord was cut off the lateral most edge and placed in 1X HBSS. The dorsal horn strips were digested in a papain solution (45 μL papain, no. 3126; Worthington Biochemical, Lakewood, NJ, United States in 3 mL 1X HBSS) for 30 minutes at 37 °C with 5% CO2, swirling every 5 minutes. The supernatant was removed and tissue was washed twice with 1 mL ice cold HEPES/HBSS solution. The tissue was then washed with 1 mL pre-warmed media consisting of Eagle’s Minimum Essential Medium with Earle’s salts and L-glutamine (10-010-CV, Corning^®^, Corning, NY, United States), 10% v/v horse serum (26050-088; Gibco, Thermo Fisher Scientific, Waltham, MA, United States), 1% v/v 100X MEM vitamin solution (11120-052; Gibco, Thermo Fisher Scientific, Waltham, MA, United States), and 1% v/v penicillin-streptomycin (Gibco, Thermo Fisher Scientific, Waltham, MA, United States). The supernatant was then removed and the tissue was triturated with 2 mL pre-warmed media using a fire-polished Pasteur pipette and centrifuged 1000 x g for 5 minutes. The supernatant was removed and the pellet was resuspended in 3 mL pre-warmed media and plated at ~1.2 × 10^5^ cells/well.

### Fluorescence Intensity Plate Readings

Fluorescence intensity plate reading (FLIPR) assays were performed on cultured DRG and sensory neuron cells 24 hours post isolation. Potassium flux of cells was measured using FLIPR Potassium Assay Kit (Molecular Devices, LLC, San Jose, CA) according to manufacturer instructions (Shieh et al., 2007). Cell culture media was replaced with a 1:1 mixture of 1X HBSS (SH30588.02, GE Healthcare Life Sciences, United States) with 20 mM HEPES (BP310-1, Thermo Fisher Scientific, Waltham, MA, United States) and FLIPR Loading Dye with 5 mM probenecid. Cells were incubated for 1 hour at room temperature in the dark before opioids were added (10 μL, in saline vehicle) and incubated for an additional 10 minutes. A nine-point dose response curve (10^−4^ M to 10^−12^ in saline) was collected in addition to saline controls. Fluorescence data was collected after the addition of 60 μL of 10 mM thallium sulfate solution per well. Background fluorescence was measured for 30 seconds in 21 second intervals before addition of opioids and after 10 minute opioid incubation. Fluorescence post-thallium addition was monitored every 21 seconds for 600 seconds total using a BioTek Synergy 2 (BioTek, Winooski, VT, United States) multi-well plate reader equipped with an excitation filter of 485/20 nm and emission filter 528/20 nm. Four technical replicates were performed for buprenorphine, DAMGO, fentanyl, and morphine in SUR1 KO and SUR1 WT mice.

### Open Field Testing

Mouse activity was measured after placement in the center of a 40 cm × 40 cm open field arena with a camcorder situated above enclosure (HDR-CX405; Sony Corp., Tokyo, Japan). Equal light distribution (45-65 lux) in the arena was verified using a digital light reader. After a 15 minute baseline recording, opioids were administered and behavior was recorded for an additional 30 minutes. Recordings were analyzed offline using Ethowatcher^®^ (developed by the Laboratory of Comparative Neurophysiology of the Federal University of Santa Catarina, freely available on www.ethowatcher.ufsc.br, IEB-UFSC (Crispim Junior et al., 2012)) which translates movement of the animal for frame-by-frame analysis. The parameters used for comparison across groups were distance traveled, time spent immobile, velocity, and total change in angular direction (in degrees).

### Adeno-Associated Virus Serotype 9 (AAV9)-Mediated SUR1 Knockdown

AAV9-mediated Cre expression in SUR1 flox mice was achieved by intrathecal injection of either AAV9-hSyn-GFP-Cre or AAV9-hSyn-GFP-ΔCre (10 μL containing ~10^13^ vector genomes, University of Minnesota Viral Vector Core, Minneapolis, MN, United States). Verification of mRNA knockdown was achieved using quantitative polymerase chain reaction and histological sections were taken from some animals in order to demonstrate successful delivery of AAV vectors by visualization of green fluorescent protein(Fisher et al., 2019).

### Data Analysis

Mechanical and thermal paw withdrawal data was analyzed for differences between genotypes using one-factor repeated measures ANOVA and two-factor repeated measures ANOVA for genotype and sex differences. In vitro fluorescence data was obtained by summing the fluorescence intensities for each opioid concentration over the 2 minute period following the addition of thallium. A dose response curve was generated and the area under the curve calculated for each technical replicate. Total fluorescence was compared between SUR1 KO and WT mice using the Mann-Whitney U test. The open field testing measurements obtained post opioid administration were separated into five-minute bins with differences between genotype and sex analyzed using repeated measures ANOVA. No significant differences were seen between male and female mice during behavioral tests, so these data were pooled. All data were analyzed using GraphPad Prism 8 (GraphPad Software, Inc., San Diego, CA), p-values < 0.05 were considered significant. Data are presented as wither mean ± SEM or as medians with 95% confidence intervals as appropriate.

## Results

### Mice lacking SUR1 subtype K_ATP_ channels have altered antinociception after buprenorphine, fentanyl, or DAMGO administration

Mice deficient in SUR1-type K_ATP_ channels display decreased morphine mechanical antinociception (Fisher et al., 2019). To compare the antinociceptive effects of synthetic and semi-synthetic opioids including buprenorphine, fentanyl, and DAMGO, mice lacking the SUR1 regulatory subunit of K_ATP_ channels (SUR1 KO) were subjected to mechanical paw withdrawal threshold testing. Administration of either buprenorphine (Figure 1A), fentanyl (Figure 1B) or DAMGO (Figure 1C) increased mechanical thresholds over the course of 60 or 120 minutes post-drug administration. Calculated area under the curve for mechanical thresholds indicate a significant difference between SUR1 WT and SUR1 KO mice after buprenorphine (unpaired t-test, p = 0.0002), fentanyl (unpaired t-test, p = 0.0018), and DAMGO (unpaired t-test, p = 0.0109) over the entire time course post-injection (Figure 1D). While mechanical nociception was attenuated with morphine administration, thermal nociception was not as significantly affected in SUR1-type K_ATP_ deficient mice (Fisher et al., 2019). A similar decrease in thermal withdrawal latency was also seen after buprenorphine administration (Figure 1E), but not after fentanyl or DAMGO injection (Figure 1F-H). Calculated area under the curve also indicated a significant decrease in TPW latencies after buprenorphine administration (Figure 1H, unpaired t-test, p = 0.0210) but not after fentanyl or DAMGO administration. The loss of K_ATP_ channel activity in the nervous system attenuates antinociception in synthetic and semi-synthetic opioids, similar to morphine.

**Figure 1.**
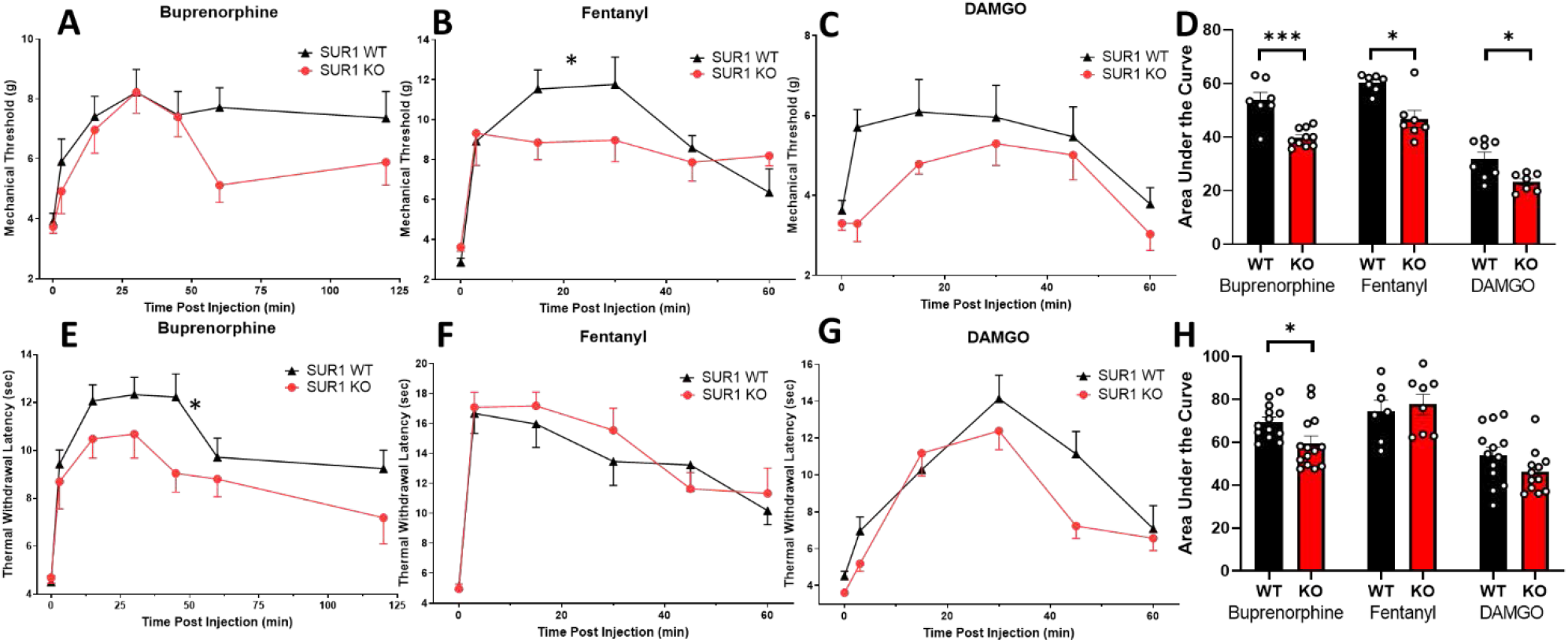
Mice lacking SUR1 subtype K_ATP_ channels have decreased antinociception to buprenorphine and DAMGO. Baseline mechanical paw withdrawal thresholds were measured for all SUR1 WT and SUR1 KO mice prior to and after administration of (A) buprenorphine (5.83 mg/kg, s.c.), (B) fentanyl (0.5 mg/kg, s.c.) and (C) DAMGO (10 mg/kg, s.c.) over 60 or 120 minutes. All three opioids increased mechanical paw withdrawal thresholds in both SUR1 WT and SUR1 KO mice. A significant interaction was found between genotypes over time after fentanyl administration (B, repeated measures ANOVA, F (5, 55) = 2.415, p = 0.0474). Area under the curve (D) for buprenorphine (unpaired t-test, p = 0.0002), fentanyl (unpaired t-test, p = 0.0018), and DAMGO (unpaired t-test, p = 0.0109) mechanical thresholds indicates a significant decrease in thresholds of SUR1 KO mice compared to SUR1 WT mice over the entire time course post-injection. Baseline thermal paw withdrawal latencies were measured for all SUR1 WT and SUR1 KO mice prior to and after administration of (E) buprenorphine (5.83 mg/kg, s.c.), (F) fentanyl (0.5 mg/kg, s.c.) and (G) DAMGO (10 mg/kg, s.c.) over 60 or 120 minutes. A significant genotype effect was found after buprenorphine administration (E, repeated measures ANOVA, F (2, 37) = 4.624, p = 0.0161). Area under the curve (H) for buprenorphine (unpaired t-test, p = 0.0210) indicates a significant decrease in latencies of SUR1 KO mice compared to SUR1 WT mice over the entire course of testing post-injection. * and *** represent significant differences (*P*<0.05 and *P*<0.001 respectively). Each curve represents the mean± SEM response of 7-13 male and female mice.

### Potassium flux is decreased in DRG and spinal cord dorsal horn cells from SUR1 KO mice after acute buprenorphine or fentanyl exposure

A global loss of SUR1-subtype K_ATP_ channels potentiates mechanical hypersensitivity in mice and attenuates morphine antinociception (Luu et al., 2019a). K_ATP_ channels are expressed in DRG and in spinal cord dorsal horn (Zoga et al., 2010), and are therefore a possible site of analgesic action of opioids. Localized deletion of the same K_ATP_ channel subtype in the lumbar dorsal horn and DRG also potentiate mechanical hypersensitivity, and decrease potassium flux in DRG after chronic morphine exposure (Fisher et al., 2019). Potassium flux was similarly examined after acute exposure to buprenorphine, fentanyl, and DAMGO in DRG and spinal cord dorsal horn isolations. DRG and spinal dorsal horn were isolated from SUR1 KO and SUR1 WT mice and these cells were exposed to ascending concentrations of either buprenorphine, fentanyl or DAMGO and fluorescence was measured over the course of 2 minutes (Figure 2). Overall, potassium flux decreased in SUR1 KO DRG following acute exposure to buprenorphine, fentanyl, and morphine (Figure 2A, Mann Whitney U, p = 0.0286). Potassium flux was also decreased in SUR1 KO spinal dorsal horn after acute buprenorphine and fentanyl administration (Figure 2B, Mann Whitney U, p = 0.0286).

**Figure 2.**
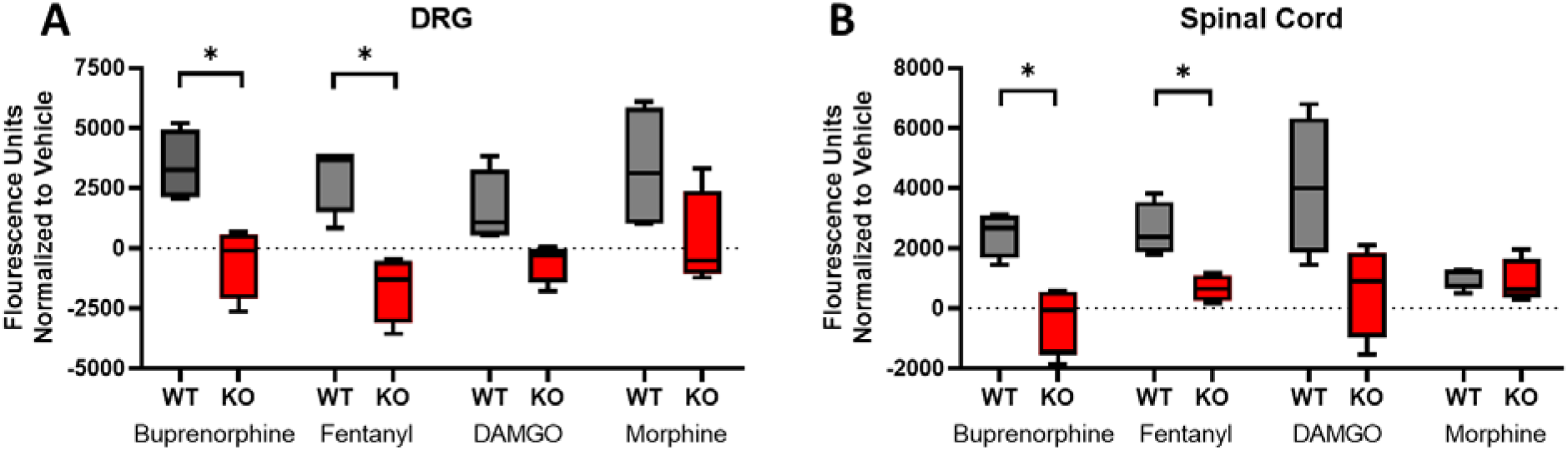
Potassium flux is decreased in DRG and spinal cord dorsal horn cells from SUR1 KO mice after acute opioid treatment. Fluorescence was pooled across a nine-point dose response curve and averaged across four trials from either SUR1 WT or KO mice. (A) Potassium flux in DRG is significantly attenuated after acute administration of buprenorphine (Mann–Whitney *U*, p = 0.0286) and fentanyl (Mann–Whitney *U*, p = 0.0286). (B) Potassium flux in spinal cord dorsal horn cells is significantly attenuated after acute administration of buprenorphine (Mann–Whitney *U*, p = 0.0286) and fentanyl (Mann–Whitney *U*, p = 0.0286). Minimum to maximum values and median values shown in the box-and-whiskers plots.

### Hyperlocomotion after acute administration of morphine and buprenorphine is potentiated in mice lacking SUR1 subtype KATP channels

Systemic administration of morphine or other drugs of abuse can lead to “drug-seeking” behaviors, including hyperlocomotion. In order to examine acute opioid-induced hyperlocomotion, open field tests examining distance traveled, time spent immobile, velocity, and total change in orientation angle were examined (Mirkovic et al., 2012). Locomotor activity has been shown to increase after morphine administration and decrease after morphine is withdrawn, and is sometimes used as a pseudo-indicator of drug-seeking activity in rodents(Shen et al., 2010; Zhang and Kong, 2017). The hypothesis that a loss of function of K_ATP_ channels would change novelty seeking behaviors was tested. After administration of 5 mg/kg morphine, the distance traveled within an open field arena was significantly higher for SUR1 KO compared to SUR1 WT mice (Figure 3A; repeated measures ANOVA, F (1, 12) = 4.957, p = 0.0459). The time spent immobile was also significantly decreased for SUR1 KO mice compared to SUR1 WT mice (Figure 3B; repeated measures ANOVA, F (1, 12) = 5.240, p = 0.0410). Animal velocity was not significantly different between the treatment groups (Figure 3C), but the change in angular orientation was significantly increased in SUR1 KO compared to SUR1 WT mice (Figure 3D, repeated measures ANOVA, F (1, 12) = 10.16, p = 0.0078). A dose of 15 mg/kg morphine resulted in similar findings including a significant increase in distance traveled (Figure 3E, repeated measures ANOVA, F (1, 28) = 7.752, p = 0.0095) and a significant decrease in time spent immobile (Figure 3F, repeated measures ANOVA, F (1, 28) = 13.11, p = 0.0012) in SUR1 KO compared to SUR1 WT mice. The speed and change in angular orientation were not significantly different across genotypes (Figure 3G-H). Spatially restricted loss of SUR1-subytpe KATP channels by intrathecal injection of AAV9-hSyn-Cre in SUR1 flox mice did not significantly change locomotion compared to control animals, indicating ascending projections do not substantially contribute to hyperlocomotion after morphine administration (Supplemental Figure 1).

**Figure 3.**
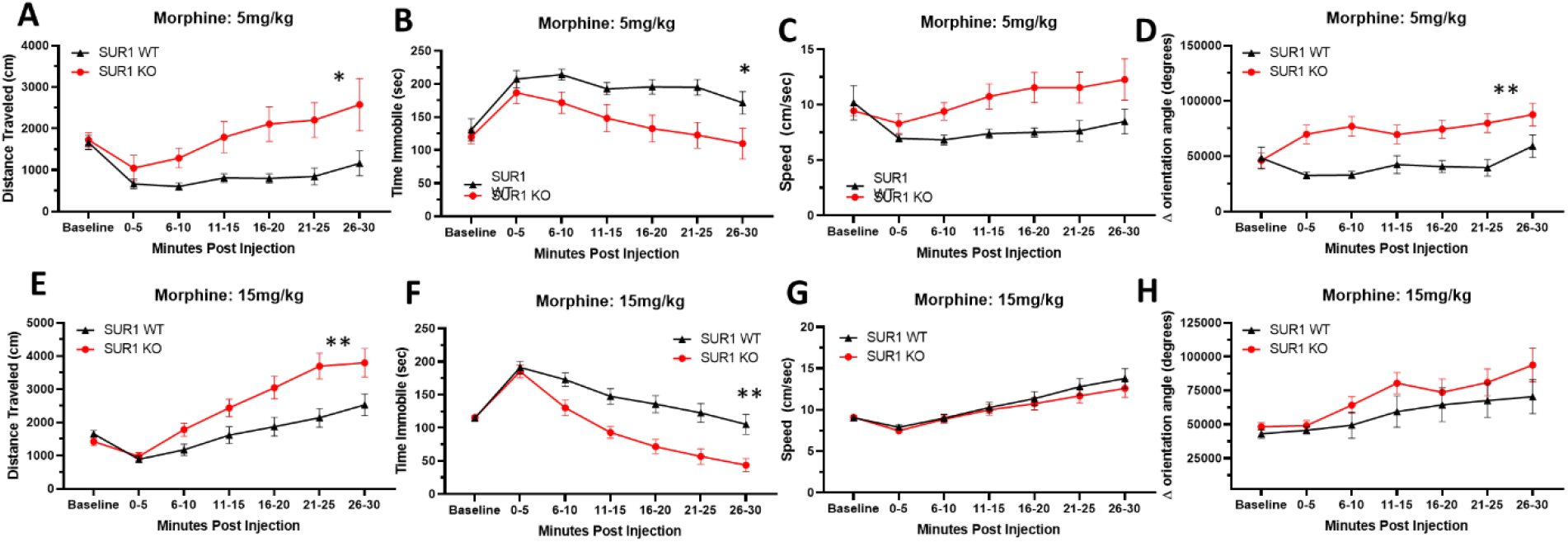
Hyperlocomotion after acute administration of morphine is potentiated in SUR1 KO mice. After a 15 minute baseline analysis, locomotion parameters were measured in mice following injection of either 5 mg/kg (A-D) or 15 mg/kg morphine (E-H). SUR1 KO mice had a significantly higher distance traveled (A) than SUR1 WT after 5mg/kg (factor: genotype, F (1, 12) = 4.957, p = 0.0459) and (E) 15 mg/kg morphine (factor: genotype, F (1, 28) = 7.752, p = 0.0095). The time spent immobile was significantly lower in SUR1 KO mice compared to SUR1 WT mice after (B) 5 mg/kg (factor: genotype, F (1, 12) = 5.240, p = 0.0410) or (F) 15 mg/kg morphine (factor: genotype, F (1, 28) = 13.11, p = 0.0012). The overall velocity (C, G) was not significantly different between SUR1 KO and SUR1 WT mice, but the change in angular orientation was significantly higher in SUR1 KO compared to SUR1 WT mice after (D) 5 mg/kg morphine (factor: genotype, F (1, 12) = 10.16, p = 0.0078) but not 15 mg/kg (H). Data analyzed by repeated measures ANOVA where * and ** represent significant differences (*P*<0.05 and *P*<0.01 respectively). Each curve represents the mean± SEM response of 5-9 male and female mice per group in A-D and 15 male and female mice per group in E-H.

In the current study and previous study, SUR1 KO mice have a loss of mechanical and thermal antinociception after acute morphine and buprenorphine administration, and hyperlocomotion was attenuated in SUR1 KO animals after morphine administration. Therefore, the ability of buprenorphine to attenuate hyperlocomotion was tested. The distance traveled after buprenorphine administration was significantly higher in SUR1 KO mice compared to SUR1 WT mice (Figure 4A; repeated measures ANOVA, interaction: F (6, 192) = 3.331, p = 0.0038). Conversely, time spent immobile was also significantly decreased in SUR1 KO mice compared to SUR1 WT mice (Figure 4B; repeated measures ANOVA, interaction: F (6, 192) = 2.387, p = 0.0301). Both animal velocity (Figure 4C) and change in angular orientation (Figure 4D) of SUR1 KO mice were significantly higher compared to SUR1 WT mice (Figure 4C; repeated measures ANOVA, F (6, 192) = 5.194, p <0.0001, Figure 4D; repeated measures ANOVA, F (6, 192) = 3.130, p = 0.006).

**Figure 4.**
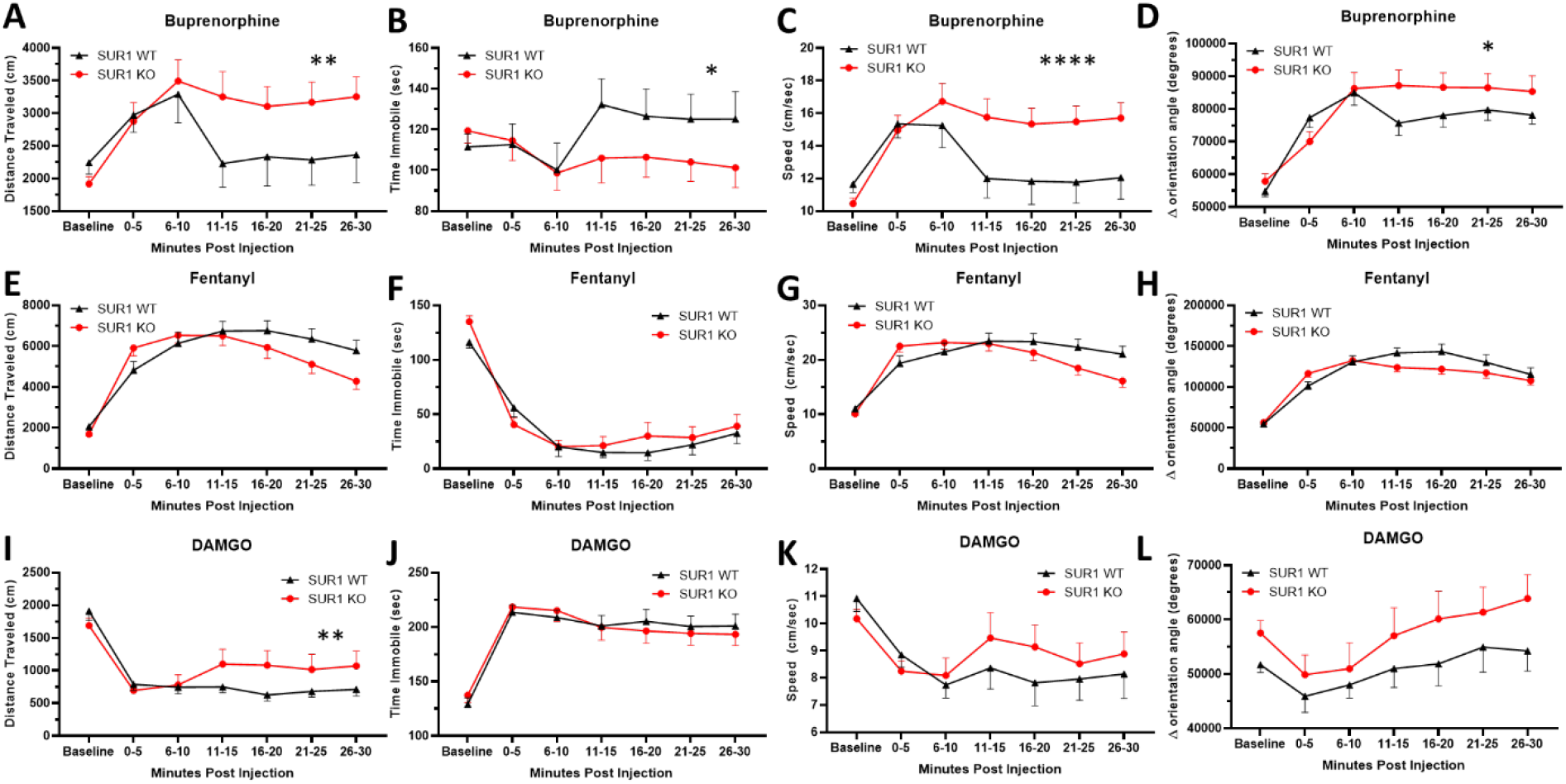
Hyperlocomotion after acute administration of buprenorphine is potentiated in SUR1 KO mice. After a 15 minute baseline analysis, locomotion parameters were measured in mice following injection with either buprenorphine (5.83 mg/kg, s.c., A-D), fentanyl (0.5 mg/kg, E-H) or DAMGO (10 mg/kg, I-L). The distance traveled in an open field (A) was significantly higher for SUR1 KO compared to SUR1 WT mice (interaction: genotype x time, F (6, 192) = 3.331, p = 0.0038) and (B) the amount of time spent immobile was significantly lower in SUR1 KO mice compared to SUR1 WT mice (interaction: genotype x time, F (6, 192) = 2.387, p = 0.0301) after buprenorphine injection. The overall velocity (C) and the (D) change in angular orientation was significantly higher in SUR1 KO compared to SUR1 WT mice after buprenorphine administration (C; interaction: genotype x time, F (6, 192) = 5.194, p <0.0001) (D; interaction: genotype x time, F (6, 192) = 3.130, p = 0.006). There was no change in locomotor activity after administration of fentanyl (E-H). SUR1 KO mice had a significantly higher (I) distance traveled (interaction: genotype x time, F (6, 198) = 2.899, p = 0.0099) compared to SUR1 WT mice after DAMGO administration. Data analyzed by repeated measures ANOVA where *, **, and **** represent significant differences (*P*<0.05, P < 0.01, and *P*<0.0001 respectively). Each curve represents the mean± SEM response of 17 male and female mice per group in A-D, 17-18 male and female mice per group in E-L.

Hyperlocomotion after administration of fentanyl was not significantly different between SUR1 KO and SUR1 WT animals (Figures 4E-H). After DAMGO treatment, the distance traveled was significantly higher for SUR1 KO mice compared to SUR1 WT mice (Figure 4I, repeated measures ANOVA, F (6, 198) = 2.899, p = 0.0099). Other measurements including time spent immobile, velocity, and change in angular orientation were not significantly different after DAMGO administration (Figures 4J-L). Overall, these data indicate loss of SUR1 subtype K_ATP_ channels affect behavioral changes attributed to opioids, in addition to their analgesic qualities.

## Discussion

In a previous study, global knockout of the SUR1 subunit of the K_ATP_ channel attenuated mechanical antinociception after morphine administration (Fisher et al., 2019). Data presented here investigated if buprenorphine, fentanyl, and the μ opioid receptor (MOR) specific agonist, DAMGO, would also produce similar effects to morphine. Two stimulus-evoked tests were used in this study to measure opioid-induced antinociception, namely the modified Hargreaves method for thermal sensitivity and von Frey method for mechanical sensitivity. This data is in agreement with previous studies finding the potentiation of morphine analgesia with concurrent use of K_ATP_ channel agonists(Afify et al., 2013; Du et al., 2011). Data presented here also demonstrate SUR1-deficiency did attenuate the antinociceptive measures of buprenorphine, DAMGO, and fentanyl with regards to mechanical sensitivity. However, thermal sensitivity did not seem to be affected to the same magnitude, concurrent with previous studies using morphine (Fisher et al., 2019). SUR1-defiency also prolonged the increase in locomotor activity after morphine and buprenorphine administration, indicating the global loss of SUR1-subtype K_ATP_ channel leads to a lack of inhibition of opioid signaling in the brain, leading to prolonged reward-seeking behavior.

Buprenorphine is a semi-synthetic derivative of an opiate alkaloid. The antinociceptive effects of buprenorphine are due to partial activation of MORs with high affinity and low intrinsic activity (Lutfy et al., 2003). Previously, buprenorphine has shown bias against MOR receptor phosphorylation and β-arrestin recruitment while super-activating adenylyl cyclase at certain doses (Boas and Villiger, 1985; Sutcliffe et al., 2017; Wang et al., 2015; Wheeler-Aceto and Cowan, 1991). Buprenorphine has a slow off rate, which helps it to displace other μ-agonists such as morphine and methadone, which is useful to treat drug dependence in the clinic. Since morphine and buprenorphine share similar chemical structures, it is not surprising the antinociceptive and hyperlocomotive actions of both drugs were attenuated in SUR1 KO mice compared to WT controls. Although buprenorphine has not been shown to induce K_ATP_ channel currents *in vitro*, the antinociceptive effects of intracerebral ventricle (icv) buprenorphine and morphine are lost after systemic K_ATP_ channel inhibition *in vivo*(Ocaña et al., 1995). It is worth noting a loss of SUR1-subtype K_ATP_ channel activity significantly modified both the antinociceptive and locomotor activities of mice after buprenorphine treatment.

The loss of SUR1 subtype K_ATP_ channels did attenuate antinociception and but did not exacerbate hyperlocomotion after fentanyl or DAMGO administration. Fentanyl is used clinically, particularly during perioperative procedures, as a potent synthetic μ receptor–stimulating opioid. The analgesic and euphoric effects of fentanyl are largely attributed to activation of the MOR, however effects are seen at alternate opioid receptors(Yoshida et al., 1999). Fentanyl’s ligand bias at MORs has been extensively characterized, and the overexpression of GPCR kinase and β-arrestin2 after fentanyl administration has been reported to enhance desensitization of u opioid-mediated GIRK potassium flux (Celver et al., 2004; Kovoor et al., 1998). However, it is unknown if fentanyl can indirectly activate or desensitize K_ATP_ channels, as seen with GIRK channels. DAMGO is also a synthetic opioid with high MOR specificity and was invented as a biologically stable analog of enkephalin. Currently DAMGO has little clinical use due to severe respiratory depression and sedation. There is almost no information on the effects of DAMGO on the activity of K_ATP_ channels in neurons, but DAMGO is reported to inhibit K_ATP_ channels in cardiomyocytes expressing subunits other than SUR1(Piao et al., 2009). These data indicate the inconsistent changes in behavioral effects after systemic administration of different opioids are through different effector mechanisms.

An open field test was used to measure locomotive behavior after opioid exposure as performed in previous studies (Banik and Kabadi, 2013; Deuis et al., 2017; Martinov et al., 2013; Mirkovic et al., 2012; Simon et al., 1994). Loss of Kir6.2 in mice produces hyperlocomotion after NMDA receptor inhibition (Zhou et al., 2012). Conversely, systemic administration of the K_ATP_ channel agonists, iptakalim, cromakalim, and pinacidil decreases amphetamine-induced hyperlocomotion(Rosenzweig-Lipson et al., 1997; Sun et al., 2010). Although the circuitry behind these results has not been thoroughly investigated, it is possible drugs of abuse facilitate activation of neuronal circuits by altering K_ATP_ channel conductance. Alteration of K_ATP_ channel activity has been shown to lead to enhanced dopamine release and activation of circuits involved in addiction (Dragicevic et al., 2015; López-Gambero et al., 2020) or indirectly through disinhibition of the ventral tegmental area through the ventromedial hypothalamus(Lee et al., 1999; van Zessen et al., 2012). Interestingly, the administration of K_ATP_ channel agonists either into the intracerebral ventricle (icv) or systemically reduces morphine withdrawal in rodents(Robles et al., 1994; Seth et al., 2010), indicating the loss of K_ATP_ channel activity may have additional roles in brain circuitry related to drug dependence. Behavioral data presented here after buprenorphine administration would support this hypothesis and previous findings with other drugs of abuse.

Behavioral data presented here in SUR1 global knockout mice after the systemic administration of structurally different opioid agonists were strikingly similar, but with some differences. Mechanically evoked thresholds were significantly attenuated in the SUR1-deficient mice for all opioids tested. This is in line with previous studies which found fentanyl analgesia could be attenuated by intraplantar administration of K_ATP_ channel antagonists(Rodrigues et al., 2005) and icv administration of the K_ATP_ channel agonist cromakalim could enhance antinociception produced by buprenorphine and morphine (Ocaña et al., 1995). Hyperlocomotion was potentiated in SUR1 KO mice after morphine and buprenorphine administration, but not after systemic delivery fentanyl or DAMGO. These differences indicate K_ATP_ channel subunits may be differentially expressed and/or modulated in areas of the nervous system, as antinociception was more affected by different classes of opioids than changes in locomotion. The degree of activation of K_ATP_ channels subtypes may be agonist and location dependent due to differences in opioid receptor coupling with the different downstream targets and protein-protein interactions(Jeske, 2019), or differences in cellular metabolism in different nervous system areas (e.g. ATP levels). We are confident that the contribution of SUR1 deficiency in the spinal cord as it relates to drug seeking behavior appears to be minimal, as intrathecal administration of AAV9-Cre viral vectors to reduce SUR1 expression in flox mice does not appear to affect locomotion significantly, but does impact morphine antinociception and tolerance(Fisher et al., 2019). Together, these data suggest K_ATP_ channel expression in the spinal cord and DRG contribute to antinociception, while K_ATP_ channel expression in higher order brain areas must be responsible for agonist-dependent drug-induced hyperlocomotion.

The buprenorphine and fentanyl induced potassium flux seen in the DRG and spinal dorsal horn was significantly attenuated in SUR1 KO mice. This is in contrast to morphine, where changes in potassium flux were only seen in the DRG of SUR1 KO mice. After DAMGO treatment, a significant decrease potassium flux was not found in either DRG or SC dorsal horn cells. This data are largely in agreement with the evoked mechanical and thermal withdrawal behavioral data indicating that a global loss of SUR1 subtype K_ATP_ channels may have direct effects on pain signaling, as SUR1 KO mice have slightly elevated mechanical thresholds compared to WT mice(Luu et al., 2019). It is also possible the administration of morphine, buprenorphine, or fentanyl also have indirect actions stimulating descending inhibitory pathways that eventually lead to K_ATP_ channel activation. Although morphine and buprenorphine are considered a prototypical μ opioid agonists, they also display activity at kappa and delta opioid receptors. Comparing the morphine and buprenorphine results with DAMGO, suggest other non-opioid mechanisms may also play a role in K_ATP_ channel-induced antinociception. Experiments using K_ATP_ channel agonists in cells or animals lacking the MOR could give an indication if these effects are exclusive to activation of one opioid receptor subtype over another, or recruitment of additional receptors.

In conclusion, the results of the present study suggest SUR1-subtype K_ATP_ channels are important in mediating the analgesic effects of several classes of opioids, while mediating some effects of drug seeking behaviors. Ongoing studies should focus on identifying the nervous system regions and neural circuits that mediate these effects in the brain and spinal cord, and if other K_ATP_ channel subtypes co-facilitate these effects.

## Contributions

Sakamaki, G; Writing - review & editing, Writing - original draft, Formal analysis, Data curation. Johnson K; Writing - review & editing, Methodology, Writing - original draft, Data curation. Mensinger M: Investigation, Data curation. Hmu E: Data curation. Klein AH; Conceptualization, Resources, Funding acquisition, Supervision, Writing - review & editing. All the authors have made a substantial contribution for the conception, design, and drafting the article. All the authors have approved the version to be submitted.

## Funding

Funding provided by the NIH to AK (K01 DA042902), the University of Minnesota Integrated Biosciences Graduate Program to GS.

## Declaration of competing interest

The authors declared no potential conflicts of interest with respect to the research, authorship, and/or publication of this article.

## Acknowledgements

The authors thank Joseph Bryan at the Pacific Northwest Diabetes Institute, Seattle, WA, United States for the SUR1 global knock out mice and SUR1 flox mice. The authors also thank Cole Fisher for her assistance with behavioral experiments.

---

**Supplemental Figure 1.**
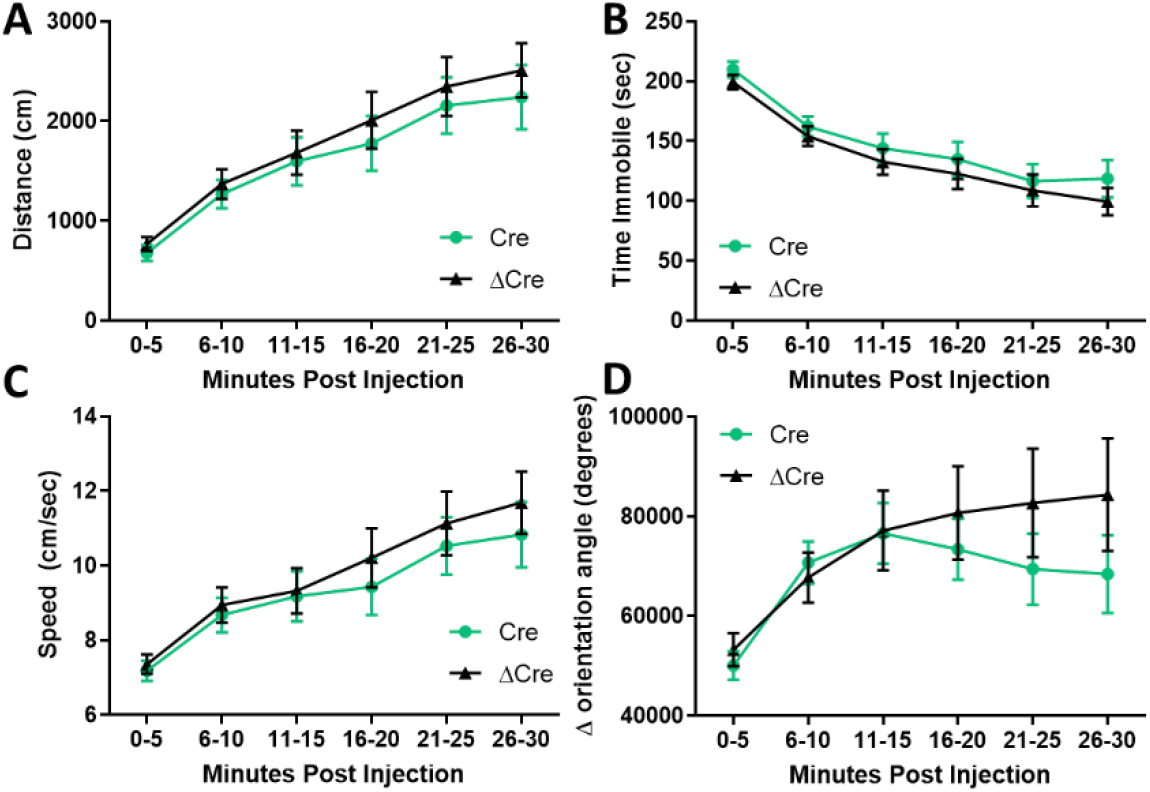
Hyperlocomotion after acute administration of morphine is not potentiated in mice with conditional deletion of SUR1. Intrathecal injection of either AAV9-hSyn-Cre or AAV9-hSyn-ΔCre does not impact (A) Distance traveled, (B) Time spent immobile, (C) Speed, or (D) the change in angular direction of mice after systemic administration of morphine (15 mg/kg, s.c.). Each curve represents the mean± SEM response of 23-24 male and female mice per group.

## Notes

### Competing Interest Statement

The authors have declared no competing interest.

## References

Afify, E.A., Khedr, M. M., Omar, A. G., Nasser, S. A., 2013. The involvement of K(ATP) channels in morphine-induced antinociception and hepatic oxidative stress in acute and inflammatory pain in rats. Fundam Clin pharmacol 27, 623–631.

Al-Hasani, R., Bruchas, M. R., 2011. Molecular mechanisms of opioid receptor-dependent signaling and behavior. Anesthesiology 115, 1363–1381.

Ashcroft, F. M., 2005. ATP-sensitive potassium channelopathies: focus on insulin secretion. The Journal of clinical investigation 115, 2047–2058.

Babenko, A. P., Aguilar-Bryan, L., Bryan, J., 1998. A VIEW OF SUR/KIR6.X, KATP CHANNELS. Annual Review of Physiology 60, 667–687.

Banik, R. K., Kabadi, R. A., 2013. A modified Hargreaves’ method for assessing threshold temperatures for heat nociception. J Neurosci Methods 219, 41–51.

Boas, R. A., Villiger, J. W., 1985. CLINICAL ACTIONS OF FENTANYL AND BUPRENORPHINE: The Significance of Receptor Binding. British Journal of Anaesthesia 57, 192–196.

Cao, D., Jing, P., Jiang, B., Gao, Y., 2017. Primary Culture of Mouse Neurons from the Spinal Cord Dorsal Horn. Bio-protocol 7, e2098.

Celver, J., Xu, M., Jin, W., Lowe, J., Chavkin, C., 2004. Distinct Domains of the μ-Opioid Receptor Control Uncoupling and Internalization. Molecular pharmacology 65, 528.

Chi, X. X., Jiang, X., Nicol, G. D., 2007. ATP-Sensitive Potassium Currents Reduce the PGE(2)-Mediated Enhancement of Excitability in Adult Rat Sensory Neurons. Brain research 1145, 28–40.

Clayton, C. C., Xu, M., Chavkin, C., 2009. Tyrosine phosphorylation of Kir3 following kappa-opioid receptor activation of p38 MAPK causes heterologous desensitization. J Biol Chem 284, 31872–31881.

Cowan, A., Doxey, J. C., Harry, E. J. R., 1977. THE ANIMAL PHARMACOLOGY OF BUPRENORPHINE, AN ORIPAVINE ANALGESIC AGENT. British Journal of Pharmacology 60, 547–554.

Crispim Junior, C. F., Pederiva, C. N., Bose, R. C., Garcia, V. A., Lino-de-Oliveira, C., Marino-Neto, J., 2012. ETHOWATCHER: validation of a tool for behavioral and video-tracking analysis in laboratory animals. Computers in Biology and Medicine 42, 257–264.

Cunha, T. M., Roman-Campos, D., Lotufo, C. M., Duarte, H. L., Souza, G. R., Verri, W. A., Funez, M. I., Dias, Q. M., Schivo, I. R., Domingues, A. C., Sachs, D., Chiavegatto, S., Teixeira, M. M., Hothersall, J. S., Cruz, J. S., Cunha, F. Q., Ferreira, S. H., 2010. Morphine peripheral analgesia depends on activation of the PI3Kgamma/AKT/nNOS/NO/KATP signaling pathway. Proc Natl Acad Sci U S A 107, 4442–4447.

Deuis, J. R., Dvorakova, L. S., Vetter, I., 2017. Methods Used to Evaluate Pain Behaviors in Rodents. Front Mol Neurosci 10, 284.

DeWire, S. M., Yamashita, D. S., Rominger, D. H., Liu, G., Cowan, C. L., Graczyk, T. M., Chen, X.-T., Pitis, P. M., Gotchev, D., Yuan, C., Koblish, M., Lark, M. W., Violin, J. D., 2013. A G Protein-Biased Ligand at the <em>μ</em>-Opioid Receptor Is Potently Analgesic with Reduced Gastrointestinal and Respiratory Dysfunction Compared with Morphine. Journal of Pharmacology and Experimental Therapeutics 344, 708.

Dragicevic, E., Schiemann, J., Liss, B., 2015. Dopamine midbrain neurons in health and Parkinson’s disease: emerging roles of voltage-gated calcium channels and ATP-sensitive potassium channels. Neuroscience 284, 798–814.

Du, X., Wang, C., Zhang, H., 2011. Activation of ATP-sensitive potassium channels antagonize nociceptive behavior and hyperexcitability of DRG neurons from rats. Mol Pain 7, 35.

Fisher, C., Johnson, K., Okerman, T., Jurgenson, T., Nickell, A., Salo, E., Moore, M., Doucette, A., Bjork, J., Klein, A. H., 2019. Morphine Efficacy, Tolerance, and Hypersensitivity Are Altered After Modulation of SUR1 Subtype KATP Channel Activity in Mice. Frontiers in Neuroscience 13, 1122.

Hill, R., Disney, A., Conibear, A., Sutcliffe, K., Dewey, W., Husbands, S., Bailey, C., Kelly, E., Henderson, G., 2018. The novel μ-opioid receptor agonist PZM21 depresses respiration and induces tolerance to antinociception. British Journal of Pharmacology 0.

Jeske, N. A., 2019. Dynamic Opioid Receptor Regulation in the Periphery. Mol Pharmacol 95, 463–467.

Kanbara, T., Nakamura, A., Shibasaki, M., Mori, T., Suzuki, T., Sakaguchi, G., Kanemasa, T., 2014. Morphine and oxycodone, but not fentanyl, exhibit antinociceptive effects mediated by G-protein inwardly rectifying potassium (GIRK) channels in an oxaliplatin-induced neuropathy rat model. Neurosci Lett 580, 119–124.

Kliewer, A., Schmiedel, F., Sianati, S., Bailey, A., Bateman, J. T., Levitt, E. S., Williams, J. T., Christie, M. J., Schulz, S., 2019. Phosphorylation-deficient G-protein-biased μ-opioid receptors improve analgesia and diminish tolerance but worsen opioid side effects. Nat Commun 10, 367.

Knapman, A., Santiago, M., Connor, M., 2014. Buprenorphine signalling is compromised at the N40D polymorphism of the human μ opioid receptor in vitro. Br J Pharmacol 171, 4273–4288.

Kovoor, A., Celver, J. P., Wu, A., Chavkin, C., 1998. Agonist Induced Homologous Desensitization of μ-Opioid Receptors Mediated by G Protein-Coupled Receptor Kinases Is Dependent on Agonist Efficacy. Molecular pharmacology 54, 704.

Lee, K., Dixon, A. K., Richardson, P. J., Pinnock, R. D., 1999. Glucose-receptive neurones in the rat ventromedial hypothalamus express KATP channels composed of Kir6.1 and SUR1 subunits. J Physiol 515 (Pt 2), 439–452.

Liu, Q., Li, Z., Ding, J.-H., Liu, S.-Y., Wu, J., Hu, G., 2006. Iptakalim inhibits nicotine-induced enhancement of extracellular dopamine and glutamate levels in the nucleus accumbens of rats. Brain Research 1085, 138–143.

Lutfy, K., Eitan, S., Bryant, C. D., Yang, Y. C., Saliminejad, N., Walwyn, W., Kieffer, B. L., Takeshima, H., Carroll, F. I., Maidment, N. T., Evans, C. J., 2003. Buprenorphine-Induced Antinociception Is Mediated by μ-Opioid Receptors and Compromised by Concomitant Activation of Opioid Receptor-Like Receptors. The Journal of Neuroscience 23, 10331.

Luu, W., Bjork, J., Salo, E., Entenmann, N., Jurgenson, T., Fisher, C., Klein, A. H., 2019a. Modulation of SUR1 K. Int J Mol Sci 20.

Luu, W., Bjork, J., Salo, E., Entenmann, N., Jurgenson, T., Fisher, C., Klein, A. H., 2019b. Modulation of SUR1 KATP Channel Subunit Activity in the Peripheral Nervous System Reduces Mechanical Hyperalgesia after Nerve Injury in Mice. International Journal of Molecular Sciences 20.

López-Gambero, A. J., Rodríguez de Fonseca, F., Suárez, J., 2020. Energy sensors in drug addiction: A potential therapeutic target. Addict Biol1, e12936.

Manglik, A., Lin, H., Aryal, D. K., McCorvy, J. D., Dengler, D., Corder, G., Levit, A., Kling, R. C., Bernat, V., Hübner, H., Huang, X.-P., Sassano, M. F., Giguère, P. M., Löber, S., Duan, D., Scherrer, G., Kobilka, B. K., Gmeiner, P., Roth, B. L., Shoichet, B. K., 2016. Structure–based discovery of opioid analgesics with reduced side effects. Nature 537, 185–190.

Martinov, T., Mack, M., Sykes, A., Chatterjea, D., 2013. Measuring changes in tactile sensitivity in the hind paw of mice using an electronic von Frey apparatus. J Vis Exp, e51212.

Martucci, C., Panerai, A. E., Sacerdote, P., 2004. Chronic fentanyl or buprenorphine infusion in the mouse: similar analgesic profile but different effects on immune responses. Pain 110, 385–392.

Mirkovic, K., Palmersheim, J., Lesage, F., Wickman, K., 2012. Behavioral characterization of mice lacking Trek channels. Frontiers in Behavioral Neuroscience 6, 60.

Nakamura, Y., Bryan, J., 2014. Targeting SUR1/Abcc8-type neuroendocrine KATP channels in pancreatic islet cells. PLoS One 9, e91525.

Niu, K., Saloman, J. L., Zhang, Y., Ro, J. Y., 2011. Sex differences in the contribution of ATP-sensitive K(+) channels in trigeminal ganglia under an acute muscle pain condition. Neuroscience 180, 344–352.

Ocaña, M., Del Pozo, E., Barrios, M., Baeyens, J. M., 1995. Subgroups among mu-opioid receptor agonists distinguished by ATP-sensitive K+ channel-acting drugs. Br J Pharmacol 114, 1296–1302.

Piao, L.-H., Wu, M., Kim, J.-H., 2009. Effects of μ-Opioid Agonist on ATP-sensitive Potassium Channel Activity in Isolated Ventricular Cardiomyocytes. Chonnam Medical Journal 45.

Robles, L. I., Barrios, M., Baeyens, J. M., 1994. ATP-sensitive K+ channel openers inhibit morphine withdrawal. Eur J Pharmacol 251, 113–115.

Rodrigues, A. R. A., Castro, M. S. A., Francischi, J. N., Perez, A. C., Duarte, I. D., 2005. Participation of ATP-sensitive K+ channels in the peripheral antinociceptive effect of fentanyl in rats. Braz J Med Biol Res 38.

Rosenzweig-Lipson, S., Thomas, S., Barrett, J. E., 1997. Attenuation of the locomotor activating effects of D-amphetamine, cocaine, and scopolamine by potassium channel modulators. Prog Neuropsychopharmacol Biol Psychiatry 21, 853–872.

Seghers, V., Nakazaki, M., DeMayo, F., Aguilar-Bryan, L., Bryan, J., 2000. Sur1 Knockout Mice: A MODEL FOR KATP CHANNEL-INDEPENDENT REGULATION OF INSULIN SECRETION. Journal of Biological Chemistry 275, 9270–9277.

Seino, S., Miki, T., 2003. Physiological and pathophysiological roles of ATP-sensitive K+ channels. Progress in Biophysics and Molecular Biology 81, 133–176.

Seth, V., Ahmad, M., Upadhyaya, P., Sharma, M., Moghe, V., 2010. Effect of potassium channel modulators on morphine withdrawal in mice. Subst Abuse 4, 61–66.

Shen, X., Purser, C., Tien, L. T., Chiu, C. T., Paul, I. A., Baker, R., Loh, H. H., Ho, I. K., Ma, T., 2010. mu-Opioid receptor knockout mice are insensitive to methamphetamine-induced behavioral sensitization. J Neurosci Res 88, 2294–2302.

Shieh, C. C., Brune, M. E., Buckner, S. A., Whiteaker, K. L., Molinari, E. J., Milicic, I. A., Fabiyi, A. C., Daza, A., Brioni, J. D., Carroll, W. A., Matsushita, K., Yamada, M., Kurachi, Y., Gopalakrishnan, M., 2007. Characterization of a novel ATP-sensitive K+ channel opener, A-251179, on urinary bladder relaxation and cystometric parameters. Br J Pharmacol 151, 467–475.

Simon, P., Dupuis, R., Costentin, J., 1994. Thigmotaxis as an index of anxiety in mice. Influence of dopaminergic transmissions. Behavioural Brain Research 61, 59–64.

Soergel, D. G., Subach, R. A., Burnham, N., Lark, M. W., James, I. E., Sadler, B. M., Skobieranda, F., Violin, J. D., Webster, L. R., 2014. Biased agonism of the μ-opioid receptor by TRV130 increases analgesia and reduces on-target adverse effects versus morphine: A randomized, double-blind, placebo-controlled, crossover study in healthy volunteers. PAIN^®^ 155, 1829–1835.

Sun, T., Zhao, C., Hu, G., Li, M., 2010. Iptakalim: a potential antipsychotic drug with novel mechanisms? Eur J Pharmacol 634, 68–76.

Sutcliffe, K. J., Henderson, G., Kelly, E., Sessions, R. B., 2017. Drug Binding Poses Relate Structure with Efficacy in the μ Opioid Receptor. Journal of Molecular Biology 429, 1840–1851.

van Zessen, R., van der Plasse, G., Adan, R. A., 2012. Contribution of the mesolimbic dopamine system in mediating the effects of leptin and ghrelin on feeding. Proc Nutr Soc 71, 435–445.

Volf, N., Hu, G., Li, M., 2012. Iptakalim Preferentially Decreases Nicotine-induced Hyperlocomotion in Phencyclidine-sensitized Rats: A Potential Dual Action against Nicotine Addiction and Psychosis. Clin Psychopharmacol Neurosci 10, 168–179.

Wang, P.-C., Ho, I.-K., Lee, C. W.-S., 2015. Buprenorphine-elicited alteration of adenylate cyclase activity in human embryonic kidney 293 cells coexpressing κ-, μ-opioid and nociceptin receptors. Journal of Cellular and Molecular Medicine 19, 2587–2596.

Wheeler-Aceto, H., Cowan, A., 1991. Buprenorphine and morphine cause antisiociception by different transduction mechanisms. European Journal of Pharmacology 195, 411–413.

Wu, X.-F., Liu, W.-T., Liu, Y.-P., Huang, Z.-J., Zhang, Y.-K., Song, X.-J., 2011a. Reopening of ATP-sensitive potassium channels reduces neuropathic pain and regulates astroglial gap junctions in the rat spinal cord. PAIN 152, 2605–2615.

Wu, X. F., Liu, W. T., Liu, Y. P., Huang, Z. J., Zhang, Y. K., Song, X. J., 2011b. Reopening of ATP-sensitive potassium channels reduces neuropathic pain and regulates astroglial gap junctions in the rat spinal cord. Pain 152, 2605–2615.

Yoshida, Y., Koide, S., Hirose, N., Takada, K., Tomiyama, K., Koshikawa, N., Cools, A. R., 1999. Fentanyl increases dopamine release in rat nucleus accumbens: involvement of mesolimbic mu- and delta-2-opioid receptors. Neuroscience 92, 1357–1365.

Zhang, J. J., Kong, Q., 2017. Locomotor activity: A distinctive index in morphine self-administration in rats. PLoS One 12, e01742721.

Zhou, Y., Liu, M. D., Fan, Y., Ding, J. H., Du, R. H., Hu, G., 2012. Enhanced MK-801-induced locomotion in Kir6.2 knockout mice. Neurosci Res 74, 195–199.

Zoga, V., Kawano, T., Liang, M. Y., Bienengraeber, M., Weihrauch, D., McCallum, B., Gemes, G., Hogan, Q., Sarantopoulos, C., 2010. KATP channel subunits in rat dorsal root ganglia: alterations by painful axotomy. Mol Pain 6, 6.

